# Dopamine modulates the integrity of the perisynaptic extracellular matrix at excitatory synapses

**DOI:** 10.1101/722454

**Authors:** Jessica Mitlöhner, Rahul Kaushik, Hartmut Niekisch, Armand Blondiaux, Christine E. Gee, Max F.K. Happel, Eckart Gundelfinger, Alexander Dityatev, Renato Frischknecht, Constanze Seidenbecher

## Abstract

In the brain, Hebbian-type and homeostatic forms of plasticity are affected by neuromodulators like dopamine (DA). Modifications of the perisynaptic extracellular matrix (ECM), controlling functions and mobility of synaptic receptors as well as diffusion of transmitters and neuromodulators in the extracellular space, are crucial for the manifestation of plasticity. Mechanistic links between synaptic activation and ECM modifications are largely unknown. Here, we report that neuromodulation via D1-type DA receptors can induce targeted ECM proteolysis specifically at excitatory synapses of rat cortical neurons via proteases ADAMTS-4 and -5. We show that receptor activation induces increased proteolysis of brevican (BC) and aggrecan, two major constituents of the adult ECM, *in vivo* and *in vitro*. ADAMTS immunoreactivity is detected near synapses, and shRNA-mediated knockdown reduced BC cleavage. We outline a molecular scenario how synaptic activity and neuromodulation are linked to ECM rearrangements via increased cAMP levels, NMDA receptor activation, and intracellular calcium signaling.

## Introduction

Synaptic transmission and plasticity are affected by perisynaptic and extrasynaptic factors including neuromodulators, glial-derived components or the extracellular matrix (ECM). The neuromodulator dopamine (DA) plays an important role in classical as well as newly discovered forms of synaptic plasticity, like neo-Hebbian or spike-timing-dependent plasticity (STDP), and hence is fundamental for various forms of learning (Tritsch and Sabatini, 2012; Lisman, 2017). Dopaminergic modulation of synapses lasts from milliseconds to hours and comprises such diverse mechanisms as regulation of presynaptic neurotransmitter release, e.g. via control of axon terminal excitability or calcium influx, postsynaptic neurotransmitter detection via regulated receptor insertion, or synaptic integration in networks [summarized in (Tritsch and Sabatini, 2012)]. Ultimately, dopaminergic activation also contributes to structural spine plasticity (Cannizzaro et al., 2019).

Dopaminergic signaling is mediated through five different G protein-coupled receptors which can be attributed to two major subgroups: D1-like and D2-like DA receptors (Seeman, 1987; Tiberi et al., 1991; Dolan et al., 1995). Both receptor subgroups have been shown to be coupled to adenylyl cyclase (AC). D1-like receptor activity leads to increased cAMP levels and activation of protein kinase A (PKA) [reviewed in (Beaulieu and Gainetdinov, 2011)]. D1 receptors (D1Rs) are localized both pre- and postsynaptically (Miklosi et al., 2018). D2 (D2R) and D3 receptors have also been found to be expressed both postsynaptically on DA target cells and presynaptically on dopaminergic neurons, respectively (Langer, 1997; Miklosi et al., 2018).

DA receptors may interact heterologously with other receptor types, such as AMPA-type (AMPARs) and NMDA-type glutamate receptors (NMDARs). This association seems to be important in regulating long-term potentiation (LTP) and working memory (Kruse et al., 2009; Nai et al., 2010; Ladepeche et al., 2013). NMDARs have been reported to form dynamic surface clusters with D1Rs in the vicinity of glutamatergic synapses. Thus, D1Rs may regulate synaptic plasticity by modulating the synaptic localization of NMDARs (Ladepeche et al., 2013).

Here, we follow the hypothesis that dopaminergic signaling can also affect the integrity of the hyaluronan (HA)-based extracellular matrix (ECM) surrounding and stabilizing synapses. This type of ECM structures occurs in the brain as highly condensed perineuronal nets (PNNs) or the more diffuse perisynaptic ECM (Dityatev et al., 2010). The perisynaptic ECM constitutes a meshwork of macromolecules based on HA as a backbone for chondroitin sulfate proteoglycans like brevican (BC), aggrecan (ACAN), versican or neurocan, glycoproteins and link proteins [for review see (Celio et al., 1998; Sonntag et al., 2015)]. As shown by us and others, *in vivo*-like ECM structures also develop in dissociated neuronal primary cultures (Miyata et al., 2005; John et al., 2006; Frischknecht et al., 2009; Faissner et al., 2010). Interestingly, the HA-based neural ECM is formed and remodeled in an activity- and plasticity-dependent manner under *in vivo* and *in vitro* conditions (Dityatev et al., 2007; Valenzuela et al., 2014). Prime candidates for this remodeling are matrix metalloproteases (MMPs) and a disintegrin and metalloprotease with thrombospondin motifs (ADAMTS) enzymes (Rauch, 2004). ADAMTS-4 was identified as one of the major proteases processing the proteoglycans ACAN, BC, neurocan and versican [reviewed in (Kelwick et al., 2015)], thus being a key candidate for ECM remodeling in the brain (Zimmermann and Dours-Zimmermann, 2008; Gundelfinger et al., 2010). ADAMTS enzymes can intrinsically be inhibited by tissue inhibitors of matrix proteases (TIMPs) and activated in response to central nervous system (CNS) injury or disease. However, the molecular pathways mediating neuronal activity-related control of ADAMTS remained elusive (Murphy, 2011; Gottschall and Howell, 2015).

The extracellular activity of the tissue plasminogen activator (tPA) protease was shown to be significantly increased in the *nucleus accumbens* (NAc) of mice after activation of D1-like DA receptors via a PKA-dependent pathway (Ito et al., 2007). This enhanced activity is probably associated with an increased release of tPA from vesicles into the extracellular space upon neuronal stimulation, what in turn regulates homeostatic and Hebbian-type synaptic plasticity (Bruno and Cuello, 2006; Ito et al., 2007; Jeanneret and Yepes, 2017). Here, we hypothesize that proteases modifying the HA-based perisynaptic ECM (i.e. ADAMTS-4 and -5) may also be released or activated after stimulation of D1-like DA receptors. Consequently, we investigated whether activation of DA receptors affects perisynaptic ECM integrity via stimulating release or activation of ECM-modifying ADAMTS proteases. We found DA receptor agonists increasing cleavage of the ECM constituents BC and ACAN in the rat cortex and in primary cortical cultures, identified the responsible matrix metalloproteases and unraveled the underlying intracellular signaling mechanisms.

## Material and Methods

### Antibodies and drugs

The following primary antibodies were used: rabbit anti-ADAMTS-4 (Abcam, Cambridge, UK), rabbit anti-ADAMTS-5 (OriGene, Rockland, MD, USA), rabbit anti-aggrecan (Merck Millipore, Burlington, MA, USA), rabbit anti-aggrecan neoepitope (Novus Biologicals, Centennial, CO, USA), guinea pig anti-brevican (Seidenbecher et al., 1995) (custom-made; central region of rat BC), mouse anti-brevican (BD Biosciences, San José, CA, USA), rabbit anti-brevican “neo” (Rb399) (custom-made; neo-epitope CGGQEAVESE) (Yuan et al., 2002; Valenzuela et al., 2014), affinity-purified rabbit anti-brevican “neo” (Rb399) (custom-made; neo-epitope CGGQEAVESE), rat anti-D1 dopamine receptor (Sigma-Aldrich, St. Louis, MO, USA), rabbit anti-D2 dopamine receptor (Abcam), mouse anti-GAD65 (Abcam), rabbit anti-GAPDH (SYSY, Göttingen, Germany), rabbit anti-GFAP (SYSY), mouse anti-Homer 1 (SYSY), mouse anti-MAP2 (Sigma-Aldrich), mouse anti-PSD95 (NeuroMab, Davis, CA, USA).

Secondary antibodies were purchased from: Cy™ 3 goat anti-rabbit IgG (H+L) (Dianova, Hamburg, Germany), Alexa Fluor® donkey anti-mouse 568 IgG (H+L) (Invitrogen, Carlsbad, CA, USA), Alexa Fluor® 488 donkey anti-mouse IgG (H+L) (Invitrogen), Alexa Fluor® 488 donkey anti-rabbit IgG (H+L) (Invitrogen), Cy™ 3 donkey anti-guinea pig IgG (H+L) (Dianova), donkey anti-mouse 647 IgG (H+L) (Invitrogen), Alexa Fluor® 488 donkey anti-rat IgG (H+L) (Invitrogen), peroxidase-coupled AffiniPure donkey anti-rabbit IgG (H+L) (Jackson ImmunoResearch, Cambridgeshire, UK), peroxidase-coupled AffiniPure donkey anti-mouse IgG (H+L) (Jackson ImmunoResearch).

Drugs were used in the given concentrations and obtained from the following companies: SKF38393 hydrobromide (for *in vivo* application 5 mg/kg body weight, Tocris Bioscience, Bristol, UK), SKF81297 hydrobromide (for *in vitro* application 1 µM, Tocris Bioscience), (-)-Quinpirole hydrochloride (1 µM, Tocris Bioscience), SCH23390 hydrochloride (10 µM, Abcam), Tetrodotoxin citrate (TTX, 1 µM, Tocris Bioscience), AP-5 (50 µM, Tocris Bioscience), Ifenprodil (3 µM, Tocris Bioscience), cAMPS-Rp triethylammonium salt (15 µM, Tocris Bioscience), recombinant human TIMP-3 (7.5 nM, R&D Systems, Minneapolis, MN, USA), Diltiazem hydrochloride (20 µM, Tocris Bioscience), KN93 phosphate (2 µM, Tocris Bioscience), Hexa-D-Arginine (0.58 µM, Tocris Bioscience).

### Primary neuronal cultures

Primary cultures of rat frontal cortical neurons were prepared for immunocytochemistry (ICC) and biochemistry as described previously (Lazarevic et al., 2011). In brief, cells were plated on poly-D-lysine-coated glass coverslips in a density of 50,000 cells per coverslip (Ø 15 mm) for ICC and in a density of 500,000 cells per poly-D-lysine-coated well (Ø 35 mm) for biochemical analysis. Cultures were kept in an incubator at 37°C and 5% CO_2_ for 21days *in vitro* (DIV).

### Immunocytochemistry

After drug treatment living neurons were incubated with antibodies against full-length and cleaved BC (Rb399) in culture medium for 20 min at 37 °C and afterwards fixed with 4 % paraformaldehyde (PFA) in phosphate-buffered saline (PBS) for 5 min at room temperature (RT). Permeabilization and blocking were done by 30 min incubation with blocking solution (10 % fetal calf serum (FCS) in PBS, 0.1 % glycine, 0.1 % Triton-X 100) at RT. Cells were incubated with primary antibodies diluted in blocking solution overnight at 4 °C. On the following day cells were washed three times with PBS for 10 min, stained with fluorescently labeled secondary antibodies for 45 min at RT in the dark, washed three times with PBS for 10 min and mounted with Mowiol (Carl Roth, Karlsruhe, Germany). Preparations were kept at 4 °C until image acquisition. Images were acquired on a Zeiss Axioplan fluorescence microscope (Carl Zeiss, Oberkochen, Germany) and further processed for quantitative analysis with NHI ImageJ 1.51w (US National Institutes of Health, Bethesda, MD, USA). Quantification of perisynaptic full-length BC and Rb399 intensities was done using OpenView Software (OpenView 1.5, Noam Ziv) (described in (Friedman et al., 2000; Shapira et al., 2003)).

### siRNA sequences

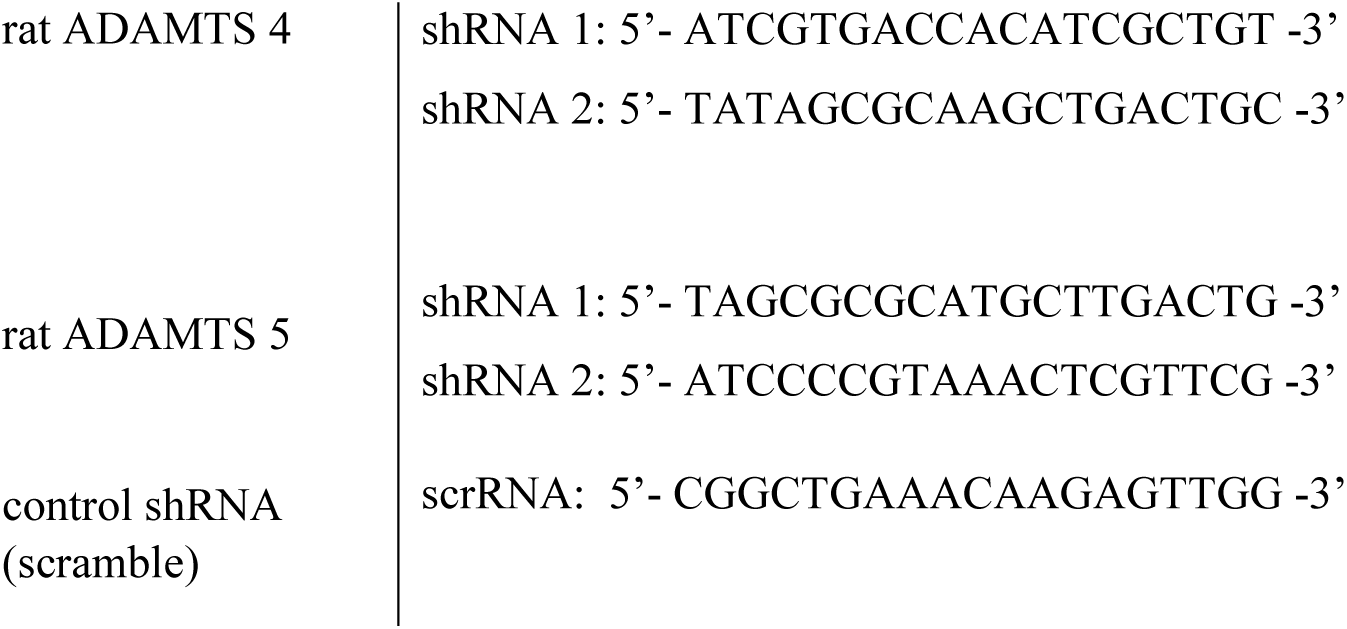

### Design of shRNA plasmids

To knockdown rat ADAMTS-4 (GeneID: 66015) and ADAMTS-5 (GeneID: 304135) shRNA plasmids were cloned by the insertion of the siRNA sequences (Dharmacon, Lafayette, CO, USA; Horizon Discovery, Waterbeach, UK) targeting the open reading frame of rat ADAMTS-4 and ADAMTS-5 into AAV U6 GFP (Cell Biolabs Inc., San Diego, CA, USA) using BamH1 (New England Biolabs, Frankfurt a.M., Germany) and EcoR1 (New England Biolabs) restriction sites.

### Generation of Adeno-associated viral particles

Positive clones were sequenced and used for the production of recombinant adeno-associated particles as described previously in (McClure et al., 2011). In brief, HEK 293T cells were transfected using calcium phosphate with an equimolar mixture of the shRNA-encoding AAV U6 GFP, pHelper (Cell Biolabs Inc.) and RapCap DJ plasmids (Cell Biolabs Inc.). 48 h after transfection, cells were lysed using freeze-thaw cycles and treated with benzonase (50 U/ml; Merck Millipore) for 1 h at 37 °C. Lysates were centrifuged at 8,000 × g at 4 °C. Afterwards, supernatants were collected and filtered using a 0.2 micron filter. Filtrates were passed through pre-equilibrated HiTrap Heparin HP affinity columns (GE HealthCare, Chicago, IL, USA), followed by washing with Wash Buffer 1 (20 mM Tris, 100 mM NaCl, pH 8.0; sterile filtered). Columns were additionally washed with Wash Buffer 2 (20 mM Tris, 250 mM NaCl, pH 8.0; sterile filtered). Viral particles were eluted using Elution Buffer (20 mM Tris, 500 mM NaCl, pH 8.0; sterile filtered). To exchange Elution Buffer with sterile PBS Amicon Ultra-4 centrifugal filters with 100,000 Da molecular weight cutoff (Merck Millipore) were used. Finally, viral particles were filtered through 0.22 µm Nalgene® syringe filter units (sterile, PSE, Sigma-Aldrich), aliquoted and stored at -80 °C.

### Knockdown of ECM-modifying proteases using shRNA

At DIV14 dissociated rat cortical cultures were infected either with shADAMTS-4, shADAMTS-5, or a scramble construct (2.07*10^7^ particles/µl). One week later, infected cells (DIV 21) were treated with SKF81297 for 15 min to stimulate D1-like DA receptors. Afterwards staining was performed as described above. However, cells were only stained for Rb399 and the synaptic marker Homer 1. Analysis and quantification were performed as indicated above. Knockdown efficiency was verified using biochemical analysis and immunocytochemical staining for either ADAMTS-4 or ADAMTS-5.

### Optogenetic modulation of cAMP in dissociated cortical neurons

To stimulate cAMP levels in dissociated rat cortical neurons, cells (DIV 14) were infected with AAV2/7.Syn-bPAC-2A-tdimer. A 500-ms flash of a 455 nm LED (0.9 mW/mm^2^) was applied to infected cultures at DIV 21. Cells were stained for the synaptic marker Homer 1 and Rb399 at different time points. BC cleavage was analyzed at Homer 1-positive synapses as described above.

### Cell lysis

For cell lysis, culture medium was aspirated and cells were washed twice with ice-cold PBS. Afterwards cells were incubated with lysis buffer (150 mM NaCl, 50 mM Tris/HCl, pH 8, 1% Triton-X 100) containing a protease inhibitor cocktail (Complete ULTRA Tablets, EDTA-free, EASYpack, Roche Diagnostics, Basel, Schweiz) for 5 min on ice. Cells were scraped off, centrifuged at 10,000 x g at 4 °C for 15 min and supernatants were prepared for SDS-PAGE.

### *In vivo* pharmacology and subcellular brain fractionation

Adult male Wistar rats were either injected with SKF38393 (5 mg/kg body weight, i.p.) or vehicle as described previously (Schicknick et al., 2008). Rats were anesthetized with isoflurane 1h after injection followed by decapitation with a guillotine. For further use the prefrontal cortex (PFC), hippocampus and rest of the brain were dissected and stored at -80 °C as described in detail elsewhere (Niekisch et al., 2019). Subcellular brain fractionation was performed according to (Seidenbecher et al., 1995). Synaptosomal fractions were harvested and incubated with Chondroitinase ABC (Sigma-Aldrich) at 37 °C for 30 min.

### SDS-PAGE and Western Blot

Samples were prepared for SDS-PAGE by adding 5x SDS loading buffer (250 mM Tris/HCl, pH 8, 50% Glycerol, 10% SDS, 0.25% bromphenol blue, 0.5 M DTT) and heating at 95°C for 10 minutes. 5-20 % Tris-glycine SDS polyacrylamide gels were run under reducing conditions. Transfer onto PVDF membranes (Merck Millipore) was performed according to standard protocols. Membranes were blocked with 5 % non-fat milk powder in TBS-T (150 mM sodium chloride, 50 mM Tris, 0.1 % (v/v) Tween20, pH 7.6) for 30 min at RT and immunodeveloped by overnight incubation at 4 °C with primary antibodies. After washing three times with TBS-T for 10 min, membranes were incubated with secondary antibodies for 60 min at RT and washed again three times with TBS-T for 10 min. Immunodetection was performed using an ECL Chemocam Imager (INTAS Science Imaging Instruments GmbH, Göttingen, Germany). Protein quantification was performed with NHI ImageJ 1.51w.

### Statistical analysis

All statistical analyses and graphical representations were performed using GraphPad Prism 5 (version 5.02, December 17, 2008). For statistical comparison between two groups, Paired- or Unpaired-sample t tests were used. Statistical comparison of multiple groups was performed by an analysis of variance (One-way ANOVA). Dunnett’s Multiple Comparison Test was used for post-hoc comparisons. Statistical tests are indicated in the figure captions. For immunocytochemical analysis four independent experiments were performed, two coverslips per condition prepared and five cells per coverslip imaged and analyzed. Here, the indicated n-number in each caption represents the number of analyzed coverslips. In the case of western blot analysis, the indicated n-number in each caption represents 4 independent experiments.

### Ethical Statement

All experimental procedures were carried out in accordance with the EU Council Directive 86/609/EEC and were approved and authorized by the local Committee for Ethics and Animal Research (Landesverwaltungsamt Halle, Germany) in accordance with the international NIH Guidelines for Animals in Research.

## Results

### Increased BC cleavage in synaptosomes after D1-like DA receptor activation *in vivo*

It has been shown previously that the performance of Mongolian gerbils in an auditory learning task is modulated via DA, as the D1/D5 DA receptor agonist SKF38393 injected systemically shortly before, shortly after or 1 day before conditioning improved learning significantly (Schicknick et al., 2008). In mice performing a similar task, we found changes in the expression levels of BC in the auditory cortex (Niekisch et al., 2019). To test if DA may be involved in changed cortical BC levels, we pharmacologically activated DA D1-like receptors using systemic application of the D1/D5 agonist SKF38393 (5 mg/kg body weight, i.p.) in rats and investigated alterations in the ECM of the PFC where D1Rs are highly expressed [reviewed in (Beaulieu and Gainetdinov, 2011)]. In addition to BC, we also included ACAN into our investigations, since this lectican is also present at perisynaptic sites (Lendvai et al., 2013), and it is expressed in the cortex, where it was shown to act as a gatekeeper for physiological plasticity (Rowlands et al., 2018). We analyzed total BC and ACAN expression levels in the homogenate and synaptosomal fractions of the PFC. For detection of proteolytically cleaved BC, we used a specific antibody against the newly emerged C-terminus (neo-epitope; Rb399) after cleavage by ADAMTS-4 or ADAMTS-5 (Nakamura et al., 2000; Hamel et al., 2005). This antibody only detects ADAMTS-4/5-cleaved fragments, but not full-length BC (Valenzuela et al., 2014). We found that the amount of full-length BC, as well as the amount of cleaved BC, is unaltered in the homogenate fraction of control and SKF38393-treated animals (Figure 1A, D, E). In the synaptosomal fraction there was a trend for reduction in the level of full-length BC by ~32 % (Figure 1A, D), while the cleaved fragment was significantly up-regulated by nearly 50 % in SKF38393-treated animals one hour after injection (Figure 1A, E). Using the neo-epitope antibody Rb399, we could confirm the increase in the amount of cleaved BC by around 50 % in the synaptosomal fraction, while levels in the homogenate were unaltered (Figure 1B, F). Accordingly, the levels of full-length and cleaved ACAN were unaltered in the homogenate fraction (Figure 1C, G, H). In the synaptosomal fraction of SKF38393-treated animals the amount of cleaved ACAN was significantly increased and levels of full-length ACAN were unaffected (Figure 1C, G, H). These data indicate that, indeed, activation of D1-like DA receptors *in vivo* modulates synapse-associated ECM components on a short time scale.

**Figure 1:**
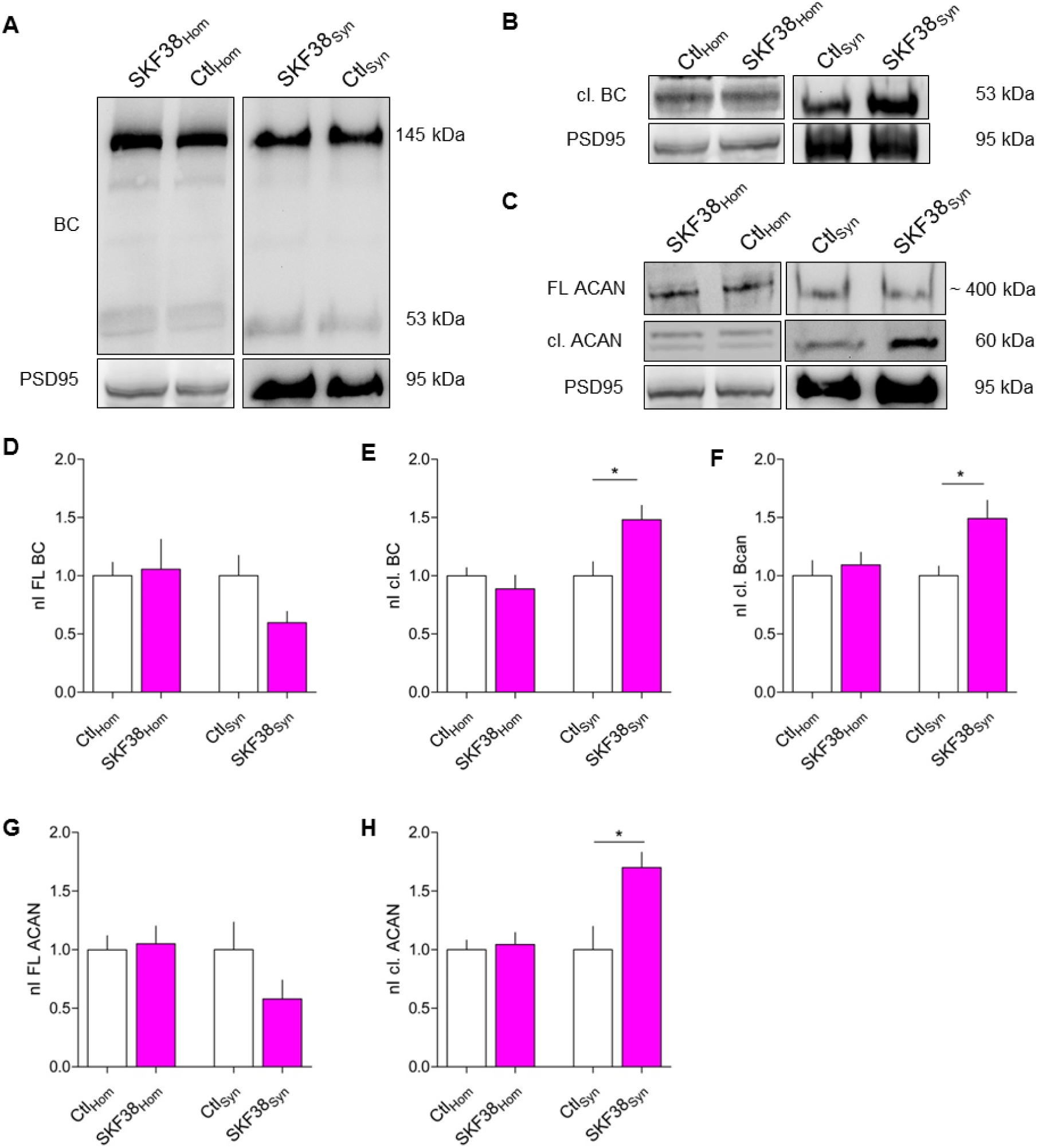
Increased cleavage of ECM lecticans BC and ACAN in synaptosomal fractions of rat PFC after systemic activation of D1-like DA receptors *in vivo*. **A)** Representitive Western blots showing the expression of BC in homogenate (left) and synaptosomal fraction (right) with and w/o SKF38393 treatment. PSD95 illustrates the efficient enrichment of synaptosomes. **B)** Representitive Western blots for the expression of cleaved BC using the neo-epitope antibody Rb399 in the homogenate (left) and the synaptosomal fraction (right). PSD95 served as an indicator for efficient fractionation. Representitive Western blots for the expression of aggrecan and the cleaved fragment at a size of 60 kDa in the homogenate (left) and the synaptosomal fraction (right). PSD95 served as an indicator for efficient fractionation. **D)** The levels of full-length BC are unalterd in the homogenate in control as well as in SKF-injected rats. In the synaptosomal fraction full-length BC is reduced after SKF38393 injection (homogenate: Ctl, 1 ± 0.1157, n = 4; SKF38, 1.056 ± 0.2574, n = 4; average ± SEM; Unpaired t test; P = 0.8491; synaptosomes: Ctl, 1 ± 0.1745, n = 4; SKF38, 0.5980 ± 0.0961, n = 4; average ± SEM; Unpaired t test; P = 0.0901) (nI FL BC = normalized intensity of full-length brevican). **E)** The expression of cleaved BC is unalterd in the homogenate after D1-like DA receptor activation, while in the synaptosomal fraction the levels of cleaved BC are significantly increased after SKF38393 treatment (homogenate: Ctl, 1 ± 0.0696, n = 4; SKF38, 0.8875 ± 0.116, n = 4; average ± SEM; Unpaired t test; P = 0.4373; synaptosomes: Ctl, 1 ± 0.12, n = 4; SKF38, 1.481 ± 0.1221, n = 4; average ± SEM; Unpaired t test; * P = 0.0308) (nI cl. BC = normalized intensity of cleaved brevican). **F)** The expression of cleaved BC as measured by the antibody Rb399 is unaltered in the homogenate but significantly increased in the synaptosomal fraction of SKF38393-injected rats (homogenate: Ctl, 1 ± 0.1301, n = 4; SKF38, 1.092 ± 0.1098, n = 4; average ± SEM; Unpaired t test; P = 0.8014; synaptosomes: Ctl, 1 ± 0.0826, n = 4; SKF38, 1.491 ± 0.1563, n = 4; average ± SEM; Unpaired t test; * P = 0.032) (nI cl. BC = normalized intensity of cleaved brevican). **G)** Full-length ACAN is unaltered in the homogenate, while it is slightly but not significant decreased in the synaptosomal fraction after D1-like DA receptor activation (homogenate: Ctl, 1 ± 0.1176, n = 4; SKF38, 1.05 ± 0.1511, n = 4; average ± SEM; Unpaired t test; P = 0.8014; synaptosomes: Ctl, 1 ± 0.2342, n = 4; SKF38, 0.5798 ± 0.1598, n = 4; average ± SEM; Unpaired t test; P = 0.1889) (nI FL Aggr = normalized intensity of full-length aggrecan). **H)** The expression of cleaved ACAN is unalterd in the homogenate, while in the synaptosomal fraction the levels are significantly increased after D1-like DA receptor activation (homogenate: Ctl, 1 ± 0.0815, n = 4; SKF38, 1.044 ± 0.1017, n = 4; average ± SEM; Unpaired t test; P = 0.7458; synaptosomes: Ctl, 1 ± 0.1985, n = 4; SKF38, 1.7 ± 0.1317, n = 4; average ± SEM; Unpaired t test; * P = 0.0261) (nI cl. Aggr = normalized intensity of cleaved aggrecan).

### D1 DA receptors are prominently expressed in rat dissociated cortical cultures

To investigate the underlying molecular mechanism of DA-regulated ECM tailoring, we used dissociated rat cortical primary neurons which express a mature type of ECM (John et al., 2006). First, we tested which types of DA receptors are expressed in the cultured neurons. For D2R, expression in rat dissociated cortical cultures has been reported previously (Sun et al., 2005). To recapitulate this finding and to prove the expression of D1R in our culture system, we performed immunocytochemical stainings at DIV 21. Indeed, both types of DA receptors appeared as little puncta along dendrites and in the vicinity of Homer 1-positive excitatory synapses (Figure 2A, B; Supplementary Figure S1A, B). While only some D1R-positive puncta were found to be present around GAD65-positive synapses (Supplementary Figure S1C), approx. 45 % of D1R-positive puncta appear in close vicinity of Homer 1-positive excitatory synapses (Figure 2C, D). A similar percentage of co-localization was found for D2R-positive puncta (Supplementary Figure S1D, E).

**Figure 2:**
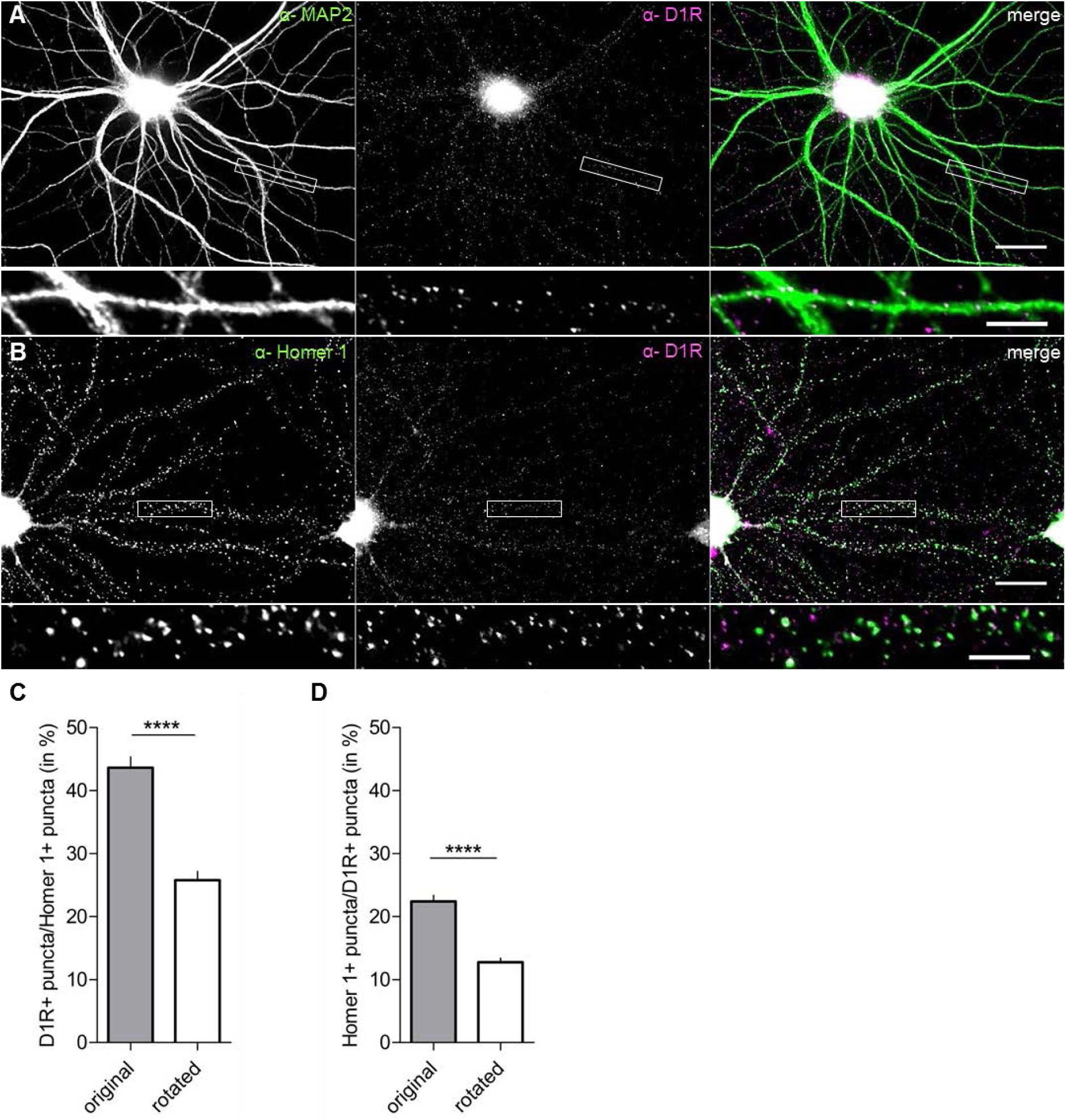
D1Rs are expressed in rat dissociated cortical neurons. **A, B)** Rat dissociated cortical neurons (DIV21) were stained for the somato-dendritic marker MAP2 (A, green) or the excitatory synaptic marker Homer 1 (B, green) and D1Rs (magenta). D1Rs appear as bright, little puncta along dendrites (A) and many of them in close vicinity of Homer 1-positive synaptic sites (B) (scale bar: 20 µm; close-up: 5 µm). **C)** In rat dissociated cortical cultures (DIV21) around 44 % of D1R-positive puncta are in close vicinity of Homer 1-positive synaptic puncta. The analysis of a 90° rotated image served as a quality control (original, 43.65 ± 1.727, n = 6; rotated, 25.79 ± 1.389, n = 6; average % ± SEM %; Paired t test; **** P < 0.0001). **D)** Around 22 % of Homer 1-positive synaptic puncta can be found in close vicinity of D1R-positive puncta. The analysis of a 90° rotated image served as a quality control (original, 22.41 ± 0.9848, n = 6; rotated, 12.75 ± 0.6574, n = 6; average % ± SEM %; Paired t test; **** P < 0.0001).

### Stimulation of D1-like, but not D2-like DA receptors augments perisynaptic BC cleavage

In a first *in vitro* approach, we analyzed BC expression and proteolytic cleavage after the stimulation of D1-like and D2-like DA receptors. To this end we treated dissociated cortical cultures at DIV 21 for 15 min with the D1/D5 DA receptor agonist SKF81297, which is commonly used for *in vitro* studies (Chen et al., 2007; Dai et al., 2008; Jürgensen et al., 2011; Li et al., 2016) or the D2-like DA receptor agonist quinpirole. Subsequently, we performed immunocytochemical stainings to quantify fluorescent signals of total and cleaved BC. As previously reported by us and others, at this time-point the cultures reliably developed a HA-based ECM with similar composition as *in vivo* (John et al., 2006; Dityatev et al., 2007; Frischknecht et al., 2009). For detection of proteolytically cleaved BC, we used the above described neo-epitope-specific antibody Rb399. The perisynaptic amount of total and cleaved BC at excitatory and inhibitory synapses was analyzed using Homer 1 and GAD65 as markers (Figure 3A, C). At Homer 1-positive excitatory synapses the amount of cleaved BC was significantly increased up to 186 ± 21% (mean ± SEM) after stimulation of D1-like DA receptors, while the amount of total BC was unaltered (Figure 3B). In contrast, at inhibitory synapses the amount of total as well as cleaved BC remained unaltered upon D1-like or D2-like DA receptor activation (Figure 3D) (data for total BC not shown). Analyzing the amount of total and cleaved BC along whole dendrites revealed no significant changes after activation of either D1-like or D2-like DA receptors (Figure 3E, F) (data for total BC not shown). Since, we observed an increase in ACAN cleavage in synaptosomes upon D1R stimulation *in vivo* (Figure 1C, H), we tested whether this also happens in the culture system. Indeed, after D1-like DA receptor stimulation the amount of perisynaptically cleaved ACAN was doubled in comparison to control conditions. Similar to BC, D2-like DA receptor activation had no effect on ACAN cleavage at Homer-positive synapses, and the amount of perisynaptic total ACAN was unaltered in both conditions (Figure 3G, H). Based on these findings we assume that DA signaling via D1-type receptors has the potential to restructure the perisynaptic ECM at excitatory synapses while D2-like receptor activation did not induce cleavage in the HA-based ECM.

**Figure 3:**
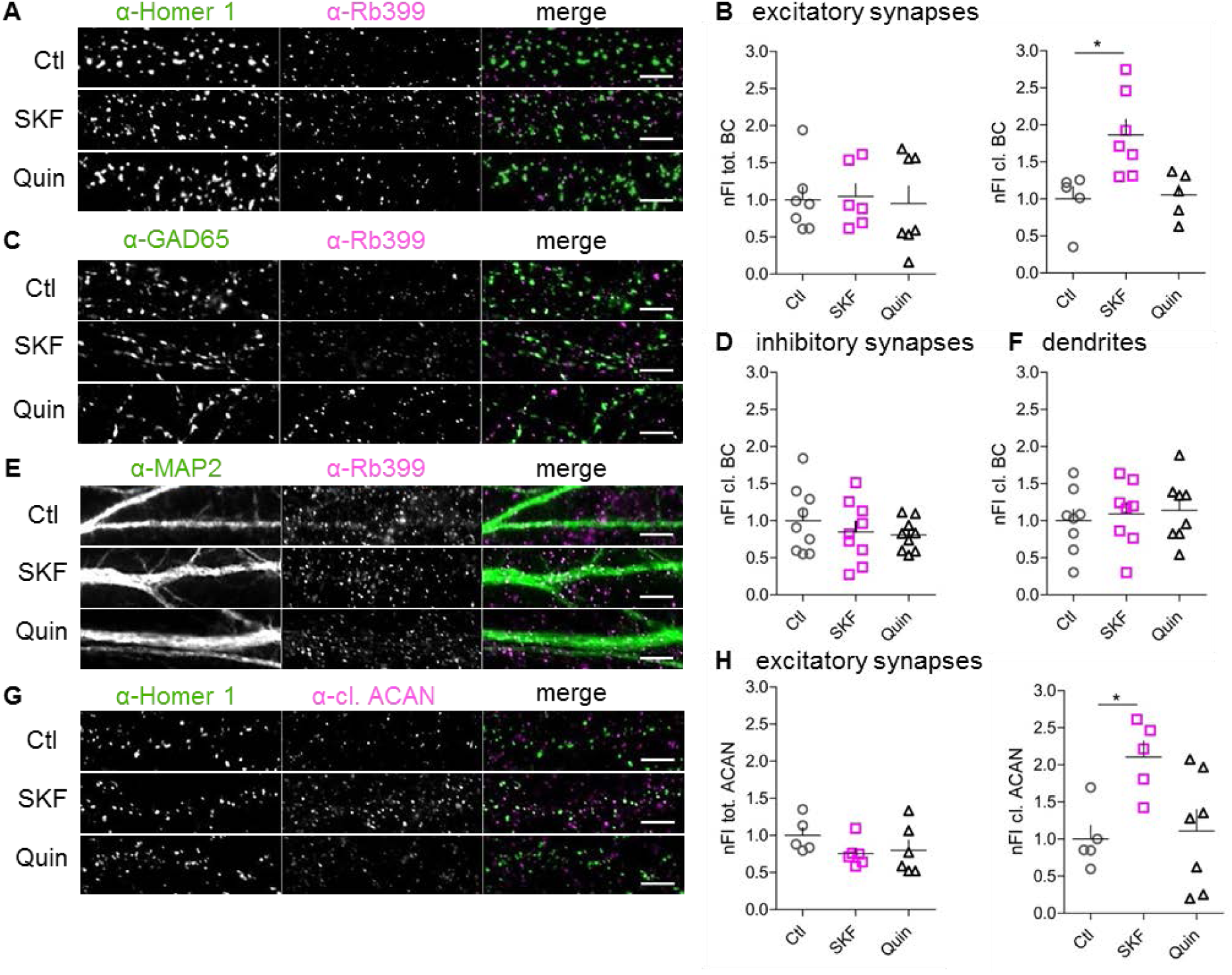
Modification of perisynaptic ECM at excitatory synapses is D1-like DA receptor-dependent. **A,C,E)** Rat dissociated cortical neurons (DIV21) were either non-treated (Ctl) or treated with SKF81297 (SKF) or quinpirole (Quin) and stained for the excitatory synaptic marker Homer 1 (A, green), the inhibitory synaptic marker GAD65 (C, green) or the somato-dendritic marker MAP2 (E, green) and extracellular cleaved BC Rb399 (magenta) (scale bar: 5 µm). **B)** At Homer 1-positive excitatory synapses the amount of cleaved BC is significantly increased after D1-like, but not after D2-like DA receptor activation (tot.BC: Ctl, 1 ± 0.1742, n = 7; SKF, 1.045 ± 0.1745, n = 6; Quin, 0.9502 ± 0.2380, n = 7; average ± SEM; One-way ANOVA; P = 0.9471; cl. BC: Ctl, 1 ± 0.1673, n = 5; SKF, 1.864 ± 0.21, n = 7; Quin, 1.054 ± 0.1403, n = 5; average ± SEM; One-way ANOVA; P = 0.007; Dunnett’s Multiple Comparison Test; * P<0.05). **D,F)** The amount of cleaved BC remained unaltered at inhibitory synapses (D) and on dendrites (F) after DA receptor activation ((D): Ctl, 1 ± 0.149, n = 9; SKF, 0.8521 ± 0.1359, n = 9; Quin, 0.8077 ± 0.0715, n = 9; average ± SEM; One-way ANOVA; P = 0.5239; (E): Ctl, 1 ± 0.1511, n = 8; SKF, 1.09 ± 0.1542, n = 8; Quin, 1.139 ± 0.1514, n = 8; average ± SEM; One-way ANOVA; P = 0.809). **G)** Rat dissociated cortical neurons (DIV21) were either non-treated (Ctl) or treated with SKF or Quin and stained for the excitatory synaptic marker Homer 1 (green) and extracellular cleaved ACAN (cl. ACAN) (magenta) (scale bar: 5 µm). **H)** ACAN cleavage was also increased at Homer 1-positive excitatory synapses after D1-like, but not D2-like DA receptor activation. The amount of total ACAN was unaltered (tot. ACAN: Ctl, 1 ± 0.1055, n = 5; SKF, 0.7565 ± 0.0728, n = 6; Quin, 0.7995 ± 0.0728, n = 6; average ± SEM; One-way ANOVA; P = 0.2938; cl. ACAN: Ctl, 1 ± 0.1859, n = 5; SKF, 2.105 ± 0.2179, n = 5; Quin, 1.108 ± 0.291, n = 7; average ± SEM; One-way ANOVA; P = 0.0204; Dunnett’s Multiple Comparison Test; * P<0.05). (nFI tot. BC = normalized fluorescent intensity of total brevican; nFI cl. BC = normalized fluorescent intensity of cleaved brevican; nFI tot. ACAN = normalized fluorescent intensity of total aggrecan; nFI cl. ACAN= normalized fluorescent intensity of cleaved aggrecan)

To prove that the augmented BC cleavage was due to activation of D1-like DA receptors, we applied the selective D1-like antagonist SCH23390 (SCH), which counteracted the SKF81297 effect at excitatory synapses (Figure 4A). The amount of cleaved BC was unaltered at inhibitory synapses and along whole dendrites after D1-like DA receptor inhibition (Figure 4B, C). Thus, perisynaptic BC cleavage at excitatory synapses, indeed, proved to be D1-like DA receptor-dependent.

**Figure 4:**
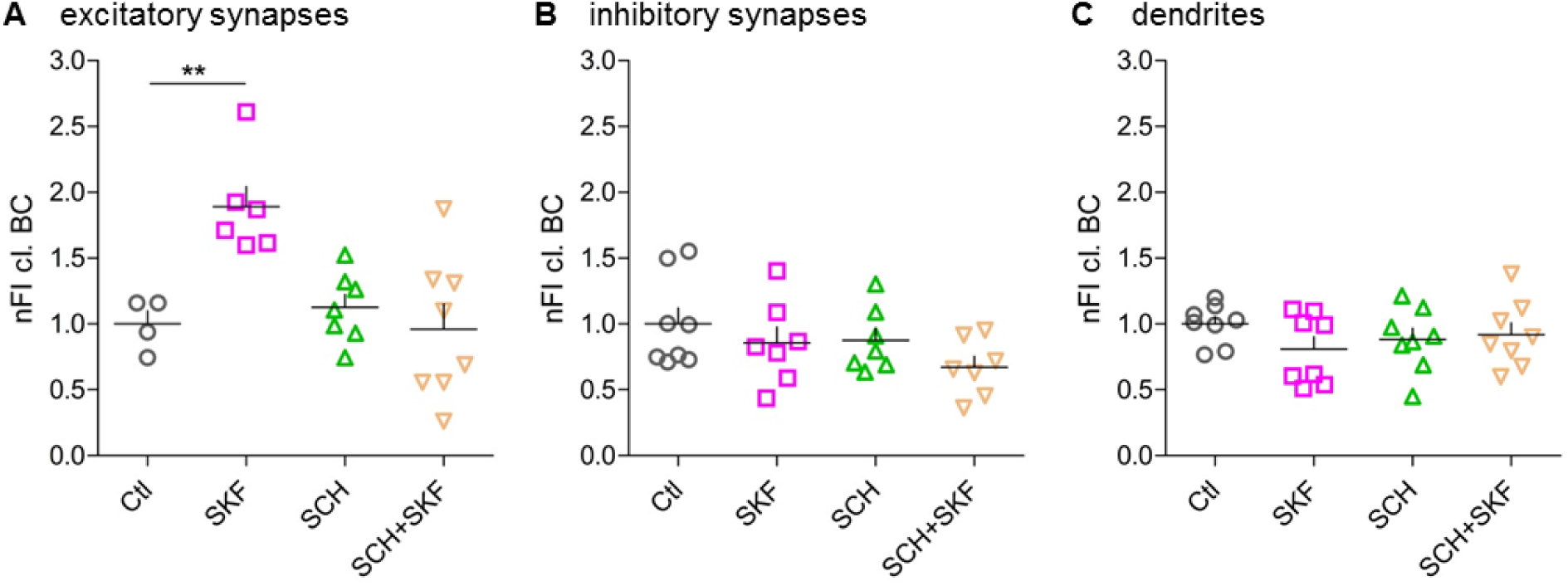
Inhibition of D1-like DA receptors suppresses SKF81297-induced BC cleavage. **A)** Treatment with SCH23390 (SCH) abolishes the SKF81297 (SKF)-induced increase in perisynaptic cleaved BC at excitatory synapses (Ctl, 1 ± 0.0999, n = 4; SKF, 1.888 ± 0.1542, n = 6; SCH, 1.125 ± 0.0998, n = 7; SKF+SCH, 0.9609 ± 0.1892, n = 8; average ± SEM; One-way ANOVA; P = 0.0014; Dunnett’s Multiple Comparison Test; ** P<0.01). **B,C)** The amount of cleaved BC was unaltered at inhibitory synapses (B) and on dendrites (C) after D1-like receptor inhibition with SCH ((B): Ctl, 1 ± 0.1217, n = 8; SKF, 0.8552 ± 0.1203, n = 7; SCH, 0.8754 ± 0.0924, n = 7; SCH+SKF, 0.6704 ± 0.0823, n = 7; average ± SEM; One-way ANOVA; P = 0.2052; (C): Ctl, 1 ± 0.054, n = 8; SKF, 0.8099 ± 0.0934, n = 8; SCH, 0.883 ± 0.0848, n = 8; SCH+SKF, 0.919 ± 0.0893, n = 8; average ± SEM; One-way ANOVA; P = 0.4384). (nFI cl. BC = normalized fluorescent intensity of cleaved brevican).

### ADAMTS-4 and ADAMTS-5 are essential for SKF81297-induced BC cleavage

Based on previous findings, the most promising candidate proteases for cleaving BC are ADAMTS-4 and ADAMTS-5 (Matthews et al., 2000; Yuan et al., 2002; Mayer et al., 2005; Viapiano et al., 2008). Therefore, we hypothesized these ECM-modifying proteases to be activated upon D1-like DA receptor stimulation. To test this, we blocked ADAMTS-4 and -5 using TIMP3, an efficient endogenous polypeptide inhibitor of these proteases. Indeed, treatment of cultures with TIMP3 alone or with TIMP3 20 min prior to D1-like receptor activation abolished the effect on BC cleavage (Figure 5A).

**Figure 5:**
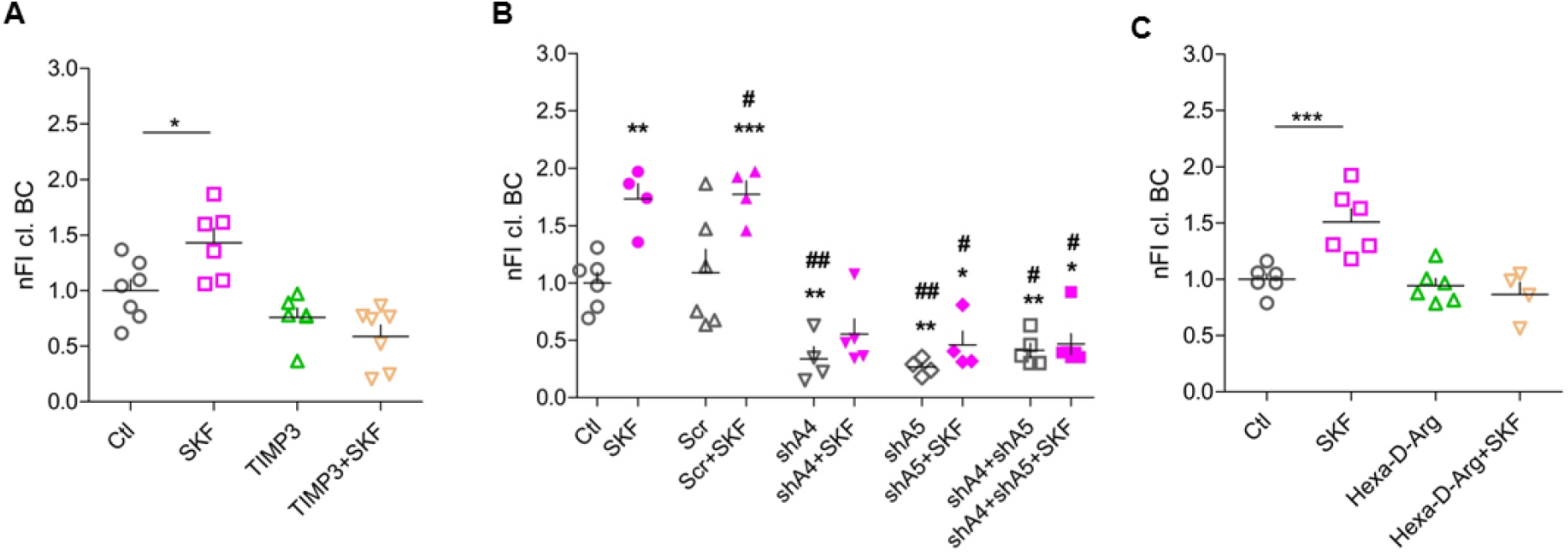
ADAMTS-4 and ADAMTS-5 are essential for D1-like DA receptor-induced BC cleavage. Inhibition of ADAMTS-4 and ADAMTS-5 with TIMP3 results in a decrease in BC cleavage around excitatory synapses (Ctl, 1 ± 0.1013, n = 7; SKF, 1.623 ± 0.2566, n = 6; TIMP3, 0.7602 ± 0.0851, n = 6; SKF+TIMP3, 0.5882 ± 0.1017, n = 7; average ± SEM; One-way ANOVA; P = 0.0489; Dunnett’s Multiple Comparison Test; * P<0.05). Knockdown of ADAMTS-4, ADAMTS-5 or both proteases together leads to a significant decrease in D1-like DA receptor-induced BC cleavage (Ctl, 1 ± 0.0929, n = 6; SKF, 1.732 ± 0.134, n = 4; Scr, 1.091 ± 0.2033, n = 6; Scr+SKF, 1.773 ± 0.1169, n = 4; shA4, 0.3385 ± 0.1052, n=4; shA4+SKF, 0.5532 ± 0.1343, n= 5; shA5, 0.2658 ± 0.0362, n=4; shA5+SKF, 0.4606 ± 0.1183, n=4; shA4+shA5, 0.4122 ± 0.0619, n=5; shA4+shA5+SKF, 0.4680 ± 0.0911, n=6; average ± SEM; One-way ANOVA; P < 0.001; Dunnett’s Multiple Comparison Test; *** P<0.001) (* = significance compared to Ctl; # = significance compared to Scr). **C)** ECM-modifying proteases are expressed in an inactive form carrying a pro-domain. Inhibition of the pro-protein convertase PACE4 diminishes SKF81927-induced BC cleavage at excitatory synapses (Ctl, 1 ± 0.051, n = 6; SKF, 1.509 ± 0.1183, n = 6; Hexa-D-Arg, 0.9441 ± 0.0636, n = 6; Hexa-D-Arg+SKF, 0.8651 ± 0.1077, n = 4; average ± SEM; One-way ANOVA; P = 0.0002; Dunnett’s Multiple Comparison Test; *** P<0.001). (nFI cl. BC = normalized fluorescent intensity of cleaved brevican)

As TIMP3 inhibits both proteases, we used a knockdown approach to differentiate between their effects. We used recombinantly produced AAVs expressing small interfering RNAs to specifically knockdown either ADAMTS-4 or -5 in our culture system. First, we tested knockdown efficiency and specificity of two different shRNA constructs for each protease using immunocytochemical stainings and Western blot analysis (data not shown). The shRNA constructs shADAMTS-4.2 (shA4) and shADAMTS-5.2 (shA5) showed the most efficient knockdown of the respective protease in neurons (Supplementary Figure S2). These constructs were used for further experiments. Next, rat dissociated cortical cultures (DIV 14) were infected with the above described shRNA constructs shA4 and shA5. At DIV 21 infected cells were treated with SKF81297 as described above and stained for Homer 1 and Rb399. As a negative control, cells were infected with AAV expressing scramble shRNA (Scr). Treatment with shRNAs shA4 and shA5 led to a strong decrease of cleaved BC under unstimulated conditions and prevented the SKF81297-induced increase in BC cleavage around excitatory synapses (Figure 5B). The concurrent knockdown of ADAMTS-4 and -5 revealed the same results as single knockdowns (Figure 5B), potentially because remaining cleaved BC is not acutely processed but stays bound at synapses. These results indicate that ECM-modifying proteases ADAMTS-4 and -5 are active under basal conditions and become more activated after D1-like DA receptor stimulation being essential for DA-dependent perisynaptic BC cleavage.

### Pro-protein furin-like convertases are involved in DA-dependent BC cleavage

ECM-modifying proteases like ADAMTS enzymes are expressed by neurons and astrocytes (Zhang et al., 2014). To become activated, their N-terminal pro-domain has to be cleaved off by pro-protein convertases (PPCs) (Tortorella et al., 2005; Kelwick et al., 2015). All members of the ADAMTS family have at least one site (R/KX_n_R/K↓R) for furin-like PPCs. To test the role of this activating step, we treated cultures with an inhibitor of furin-like PPCs, i.e. Hexa-D-Arginine. Inhibition of PPCs activities indeed abolished the SKF81297-induced increase in perisynaptic BC cleavage, suggesting DA-dependent activation or release of PPCs is upstream of ADAMTS-4/5 activation (Figure 5C).

### Network activity, activation of postsynaptic NMDARs and opening of L-type Ca^2+^ channels are essential for D1-like DA receptor-induced BC cleavage

To address the question, whether D1-like DA receptor activation is sufficient to induce increased perisynaptic ECM cleavage, we studied the role of general neuronal activity in this process. By blocking voltage-gated sodium channels using TTX to prevent action potentials and thus network activity the effect of the D1-like DA receptor agonist on BC cleavage was suppressed (Figure 6A). This suggests that neuronal activity is crucial for DA-mediated BC cleavage.

**Figure 6:**
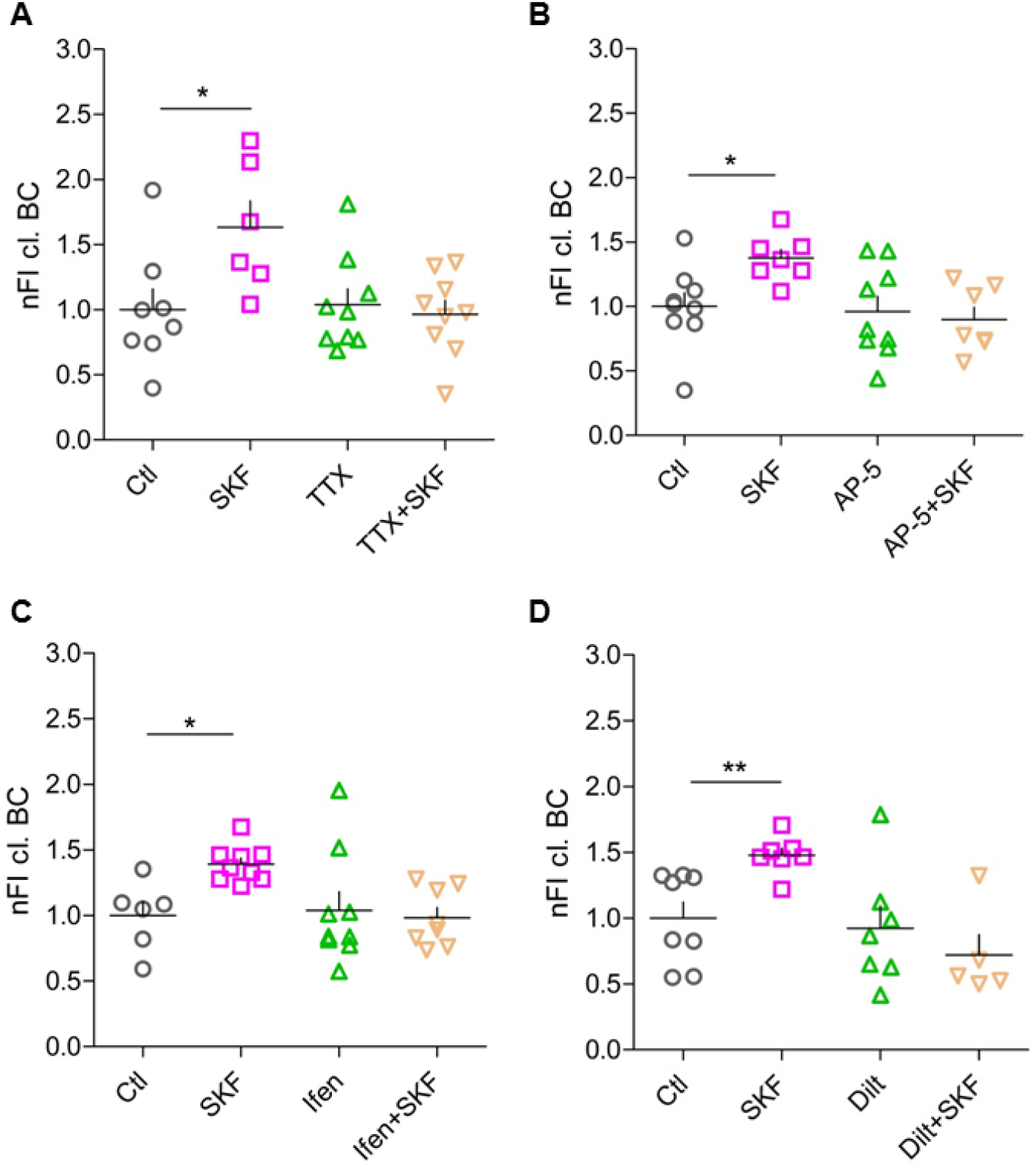
Entire network activity as well as activity of postsynaptic sites are essential for SKF81297-induced BC cleavage. **A)** Silencing neuronal networks with TTX results in an unaltered BC cleavage (Ctl, 1 ± 0.1599, n = 8; SKF, 1.632 ± 0.2036, n = 6; TTX, 1.039 ± 0.1211, n = 9; SKF+TTX, 0.9663 ± 0.1061, n = 9; average ± SEM; One-way ANOVA; P=0.0168; Dunnetti’s Multiple Comparison Test; * P<0.05). **B)** Inhibition of all types of NMDARs with AP-5 abolishes SKF81297-induced perisynaptic BC cleavage (Ctl, 1 ± 0.105, n = 9; SKF, 1.376 ± 0.0671, n = 7; AP-5, 0.9599 ± 0.1183, n = 9; SKF+AP-5, 0.8976 ± 0.0964, n = 7; average ± SEM; One-way ANOVA; P = 0.0187; Dunnett’s Multiple Comparison Test; * P<0.05). **C)** Especially, inhibition of NR2B-containing NMDARs with Ifenprodil (Ifen) results in unaltered BC cleavage (Ctl, 1 ± 0.107, n = 6; SKF, 1.392 ± 0.046, n = 9; Ifen, 1.039 ± 0.1436, n = 9; SKF+Ifen, 0.984 ± 0.0782, n = 8; average ± SEM; One-way ANOVA; P = 0.0199; Dunnett’s Multiple Comparison Test; * P<0.05). **D)** SKF81297-induced perisynaptic BC cleavage depends on signaling via L-type VGCC (Ctl, 1 ± 0.1223, n = 8; SKF, 1.478 ± 0.0542, n = 7; Dilt, 0.9228 ± 0.1698, n = 7; Dilt + SKF, 0.7188 ± 0.1541, n = 5; average ± SEM; One-way ANOVA; P = 0.0046; Dunnett’s Multiple Comparison Test; ** P<0.01). (nFI cl. BC = normalized fluorescent intensity of cleaved brevican)

Since we found changes in cleaved BC particularly around excitatory synapses, where NMDARs are located and these receptors have been shown to interact functionally with D1-like DA receptors (Haddad, 2005; Missale et al., 2006), we tested for a role of NMDARs in the signaling mechanisms underlying DA-dependent BC cleavage. To address this, we first blocked all subtypes of NMDARs using AP-5 and, indeed, abolished BC cleavage increase with this treatment (Figure 6B). Since it has been shown that D1Rs are expressed in close vicinity of GluN2B-containing NMDARs in the PFC (Hu et al., 2010), we tested if this specific receptor subtype might be a player in the pathway. Therefore, we blocked GluN2B in the culture system using Ifenprodil, which again abolished the increase in BC cleavage after D1-like receptor stimulation (Figure 6C). In summary, general network activity as well as functional postsynaptic GluN2B-containing receptors are essential to increase perisynaptic BC cleavage via DA stimulation.

Reported physiological interactions of D1-like DA receptors and L-type voltage-gated calcium channels (VGCC) suggested a possible influence of these channels in the investigated DA-dependent perisynaptic BC cleavage (Surmeier et al., 1995; Young and Yang, 2004; Chen et al., 2007). When we blocked postsynaptic L-type VGCC with the antagonist diltiazem hydrochloride, stimulation of D1-like DA receptors did not augment perisynaptic BC cleavage anymore (Figure 6D).

### Elevated intracellular cAMP levels entail enhanced BC cleavage

D1-like DA receptors are typically coupled to Gα_s/olf_ and cause increased cAMP levels and PKA activity [reviewed in (Beaulieu and Gainetdinov, 2011)]. Therefore, we investigated if elevation of intracellular cAMP levels in neurons is sufficient to result in enhanced perisynaptic BC cleavage. To do so, we used blue light-inducible AC encoded by AAV2/7.Syn-bPAC-2A-tdimer for optogenetic modulation of cAMP levels in the primary culture system as previously described (Stierl et al., 2011). BC cleavage was analyzed around Homer 1-positive synapses at different time points after blue light illumination of the cells. Already 5 min after light stimulation we observed enhanced perisynaptic BC cleavage up to nearly 150 % (Figure 7A). This effect strength is comparable to our findings of pharmacologically induced perisynaptic BC cleavage, which was found 15 min after D1-like DA receptor activation. In the optogenetic approach, the rise in intracellular cAMP levels occurs much faster. As shown by (Stierl et al., 2011), cAMP rises within about 1 second and is back to baseline about 1 min after light off in hippocampal neurons. The optogenetic cAMP elevation was sufficient to cause a brief increase in perisynaptic BC cleavage, since already 10 min after light stimulation the levels went back to baseline (Figure 7A), potentially due to highly active phosphodiesterases, which were not blocked in our experiments.

**Figure 7:**
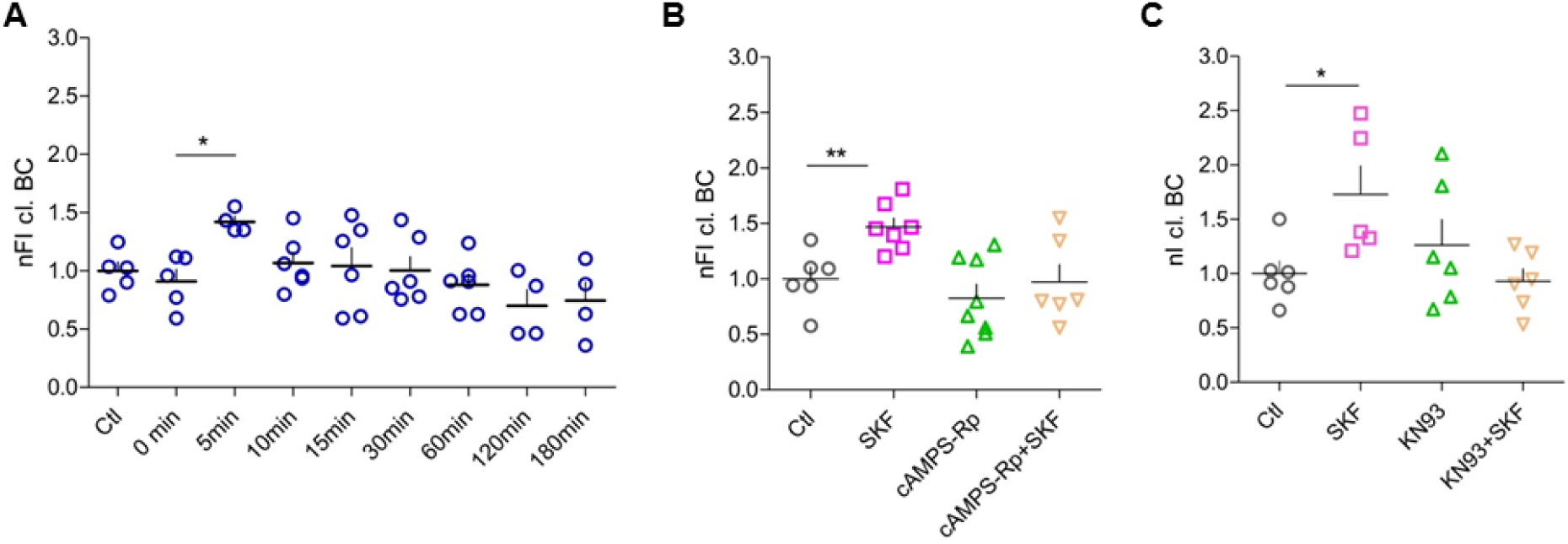
SKF81297-induced BC cleavage requires co-signaling through PKA and CaMKII. **A)** Elevated intracellular cAMP levels are crucial for an increase in BC cleavage around excitatory synapses (Ctl, 1 ± 0.0765, n = 5; 0 min, 0.9096 ± 0.1017, n = 7; 5 min, 1.42 ± 0.0481, n = 5; 10 min, 1.067 ± 0.0943, n = 6; 15 min, 1.042 ± 0.1552, n = 6; 30 min, 1.003 ± 0.1172, n = 6; 60 min, 0.8787 ± 0.0938, n = 6; 120 min, 0.6993 ± 0.1398, n = 4; 180 min, 0.7450 ± 0.1605, n = 4; average ± SEM; One-way ANOVA; P = 0.0207; Dunnett’s Multiple Comparison Test; * P<0.05). **B)** Increased perisynaptic BC cleavage after D1-like DA receptor stimulation depends on PKA activation (Ctl, 1 ± 0.1044, n = 6; SKF, 1.468 ± 0.0805, n = 7; cAMPS-Rp, 0.8243 ± 0.1249, n = 8; cAMPs-Rp + SKF, 0.9714 ± 0.1567, n = 6; average ± SEM; One-way ANOVA; P = 0.0041; Dunnett’s Multiple Comparison Test; ** P<0.01). **C)** SKF81297-induced perisynaptic BC cleavage is subjected to intracellular Ca^2+^ signaling (Ctl, 1 ± 0.1139, n = 6; SKF, 1.648 ± 0.2978, n = 5; KN93, 0.9165 ± 0.1118, n = 4; SKF+KN93, 0.9303 ± 0.1126, n = 6; average ± SEM; One-way ANOVA; P = 0.0269; Dunnett’s Multiple Comparison Test; * P<0.05). (nFI cl. BC = normalized fluorescent intensity of cleaved brevican)

### DA-dependent BC cleavage requires co-signaling through PKA and CaMKII

Based upon this reductionistic approach to show involvement of cAMP, we further characterized the intracellular signaling cascade. Accordingly, we blocked PKA activity using the cell-permeable competitive antagonist cAMPS-Rp interacting with the cAMP binding sites on the regulatory subunits of PKA and analyzed again the perisynaptic amount of cleaved BC. PKA inhibition indeed abolished the SKF81297-induced increase in perisynaptic cleaved BC confirming the hypothesis that stimulation of D1-like DA receptors results in activation/release of the extracellular proteases in a PKA-dependent manner (Figure 7B). As D1R signaling was also reported to be alternatively mediated via protein kinase C (PKC) (Lee et al., 2004; Chen et al., 2007) we also tested for the involvement of this kinase utilizing bisindolylmaleimide II (1µM), however, no effect on DA-dependent perisynaptic BC cleavage was observed (data not shown).

As BC cleavage depends on signaling through NMDARs and L-type VGCC activation indicating increased Ca^2+^ influx into the activated cells, we next tested the involvement of Ca^2+^/calmodulin-dependent kinase II (CaMKII) as a major downstream effector of Ca^2+^ signaling. Blocking CaMKII using KN93 abolished the SKF81297-induced effect of increased perisynaptic BC cleavage (Figure 7C), suggesting that co-signaling via PKA and CaMKII is required to promote BC proteolysis (Figure 8).

**Figure 8:**
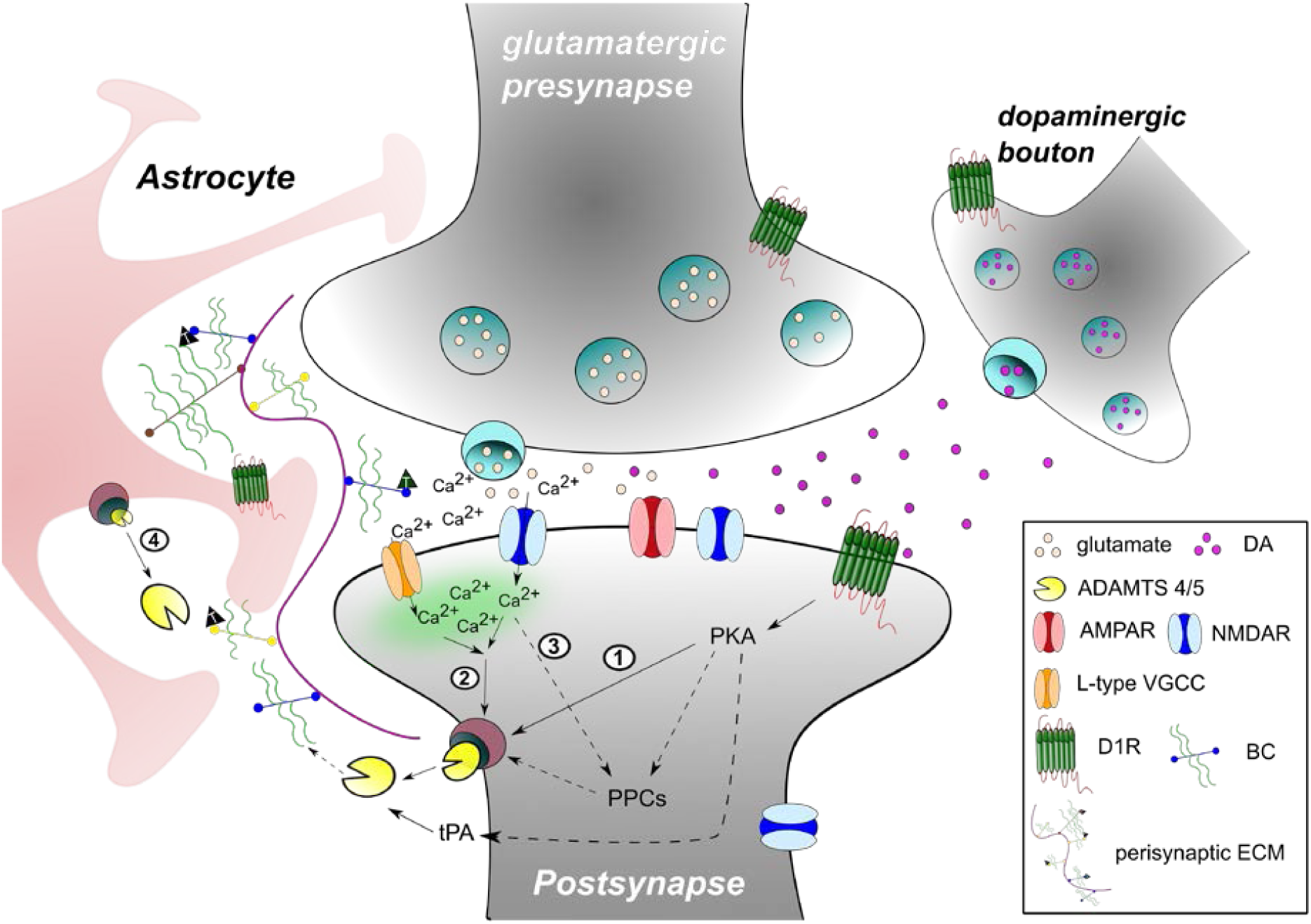
Schematic illustration of the molecular mechanisms resulting in ECM remodeling after D1-like DA receptor activation. Activation of D1 DA receptors results in increased intracellular cAMP levels leading to downstream activation of PKA. Further, active PKA might be able to release already active proteases, such as ADAMTS-4 and ADAMTS-5 in the extracellular space to remodel the perisynaptic ECM (①). Besides its downstream signaling, stimulation of D1-like DA receptor can influence the Ca^2+^ influx through NMDARs and L-type VGCC. Enhanced Ca^2+^ influx activates CaMKII that might lead to the release of ADAMTS-4 and -5 into the extracellular space (②). D1-like DA receptor activation might result in a co-signaling of pathways ① and ②. All ADAMTS enzymes carry a pro-domain to keep them in an inactive state. After D1-like DA receptor stimulation, PKA activation and enhanced Ca^2+^ influx could result in the activation of pro-protein convertases (PPCs). Active PPCs themselves cleave off the pro-domain of ADAMTS-4 and -5 resulting in active enzymes ready for release (③). ADAMTS-4 and ADAMTS-5 are expressed also by astrocytes. Therefore, activation of neuronal or astrocytic D1-like DA receptors might cause a release of these enzymes by astrocytes as well (④).

## Discussion

Here, we demonstrate that D1-like DA receptor stimulation leads to a restructuring of the ECM surrounding excitatory synapses of cortical neurons, as measured by increased BC and ACAN cleavage. This effect is mediated via the PKA-cAMP-pathway and requires network activity and NR2B/CaMKII activation, triggering extracellular proteases ADAMTS-4 and -5 (Figure 8). Our findings show that the neuromodulator DA is a potential sculptor of the perisynaptic ECM, translating dopaminergic signals into structural plasticity of excitatory synapses. These findings pave the way to a deeper mechanistic understanding of the dopaminergic contribution to neuronal and synaptic plasticity via ECM remodeling.

### How does DA contribute to local plasticity?

DA was shown to exert important neuromodulatory control over different forms of plasticity like LTP (Krug et al., 1983), long-term depression (LTD) (Calabresi et al., 1992) or STDP (Yang and Dani, 2014) [for summary see (Tritsch and Sabatini, 2012)]. In neurons of the PFC, a key structure for abstract memory and decision making, DA was shown to act both as a tonic background regulator and as a phasically released modulator of LTP or LTD, in both paradigms following inverted U-functions (Otani et al., 2015). The authors report LTP to be facilitated by tonic background DA levels, while LTD appeared to require acutely released DA for its facilitation.

Dopaminergic mechanisms were shown to be involved in spine restructuring. For instance, DA increase reverts aberrant structural plasticity in the NAc of alcohol-withdrawn rats by renormalizing the density of long thin spines and restoring limbic memory (Cannizzaro et al., 2019). In a very elegant study, Yagishita et al. (2014) demonstrated that the DA-mediated support of spine enlargement upon stimulation has to occur within a narrow time window after NMDAR activation (Yagishita et al., 2014). This mechanism of DA action involved D1Rs, PKA, NMDAR, and CaMKII, which is in good agreement with our findings presented here, thus pointing to the hypothesis that the DA-induced ECM remodelling observed by us may be involved in catecholaminergic spine morphology regulation. This is furthermore in good agreement with unpublished findings from our group showing that BC cleavage is increased in the cortex of DAT-Cre mice expressing heterozygously the Cre recombinase instead of the dopamine transporter (DAT) under the control of the DAT promoter, potentially due to chronically increased tonic DA levels because of a decrease in the maximum DA uptake rate in these animals [cf. (O’Neill et al., 2017)].

In the PFC as well as in cortical cultures, D1Rs are considered the most prominent DA receptor subtypes, but D2Rs were also found in cortical areas [(Gao and Wolf, 2007); reviewed in (Beaulieu and Gainetdinov, 2011)] and in our *in vitro* culture system. However, D2R agonists had no effect on perisynaptic ECM proteolysis. Still, the possibility exists that D1R/D2R heterodimers are involved. Since there is evidence that these heteromers are preferentially coupled to Gα_q_ G proteins, which activate PLC-IP3-DAG signaling [reviewed in (Tritsch and Sabatini, 2012)] and we found PKC not involved in the D1R-agonist-induced ECM restructuring, it is rather unlikely that D1/D2 heteromers play a crucial role, although at this point we cannot completely rule out this possibility.

Ito and colleagues (2007) identified the D1R-cAMP-PKA pathway as the underlying mechanism for enhanced extracellular tPA activity (Ito et al., 2007). Based on our findings, the perisynaptic BC cleavage upon D1-like DA receptor activation also follows this signaling pathway. Furthermore, Iwakura et al. (2011) showed in striatal cultured neurons DA- and D1R-agonist SKF38393-increased shedding and release of the epidermal growth factor (EGF) ectodomain, concomitant with ADAMTS activation and calcium signaling (Iwakura et al., 2011). Along this line, Li et al. (2016) demonstrated in striatal slices and in cultured astrocytes that the D1R-agonist SKF81297 mediated MMP activation potentiating ß-dystroglycan cleavage and leading to stimulated NMDAR calcium currents (Li et al., 2016). Taken together, the signaling pathway identified in this study (Figure 8) could be of general significance for transducing DA signals into structural and functional consequences at synapses.

### Induced ECM cleavage as a prerequisite for local structural and functional plasticity?

Restructuring or disintegration of the synaptic ECM has diverse effects on synaptic properties and characteristics, but most of the studies used exogenous glycosidase treatment rather than controlled endogenous proteolysis. Thus, glycosidic cleavage of ECM components proved to modulate synaptic short-term plasticity through exchange of postsynaptic glutamate receptors (Frischknecht et al., 2009). Further adding to this picture, synaptic LTP was also shown to be impaired after hyaluronidase treatment via involvement of VGCC (Kochlamazashvili et al., 2010). Pyka et al. (2011) showed that glycosidic cleavage of HA and chondroitin sulfate leads to an increased number of synaptic puncta and reduced postsynaptic responses in cultured hippocampal neurons (Pyka et al., 2011).

Beyond glycosidase treatment, regulated and limited proteolytic cleavage by extracellular proteases, like tPA, MMP-9, neurotrypsin or ADAM10 was also shown by some studies to affect synaptic function and structure [summarized in (Sonderegger and Matsumoto-Miyai, 2014)]. For instance, the extracellular ADAMTS protease *gon-1* was demonstrated to regulate synapse formation during development in *Caenorhabditis elegans* neuromuscular junctions (Kurshan et al., 2014).

Thus our findings presented here point to the possibility that DA-induced cleavage of perisynaptic ECM components via ADAMTS-4 and/or -5 may directly be involved in morphological restructuring of excitatory synapses. This hypothesis is consistent with studies in BC knockout mice and in rat slices treated with anti-BC antibodies showing reduced synaptic plasticity (Brakebusch et al., 2002).

### Does protease activation occur in a compartment-specific way?

Enzymes of the ADAMTS family are synthesized as pro-enzymes, which are cleaved at their C- or N-termini by furin or furin-like PPCs, such as PACE4, resulting in mature and potentially active enzymes (Flannery et al., 2002; Lemarchant et al., 2013). ADAMTS-5 was shown to be exclusively activated extracellularly (Longpré et al., 2009). However, local specificity of ADAMTS-4 activation is controversial: while one study claimed the activation of ADAMTS-4 in the *trans*-Golgi network via furin (Wang et al., 2004), Tortorella and colleagues (2005) demonstrated that at neutral pH furin is less efficient in activating pro-ADAMTS-4 than PACE4, and at acidic pH PACE4 is the only PPC efficiently activating ADAMTS-4 (Tortorella et al., 2005). Here, we demonstrated that furin or furin-like PPCs mediate DA-dependent perisynaptic BC cleavage, which is not happening at dendrites or inhibitory synapses, arguing for a locally restricted activation mechanism.

A recent study identified tPA as another potential activator for ADAMTS-4 in spinal cord injury leading to cleavage of inhibitory chondroitin sulfate proteoglycans (CSPGs) (Lemarchant et al., 2014). Interestingly, tPA has been shown to display enhanced extracellular activity depending on the activation of postsynaptic D1Rs in the NAc (Ito et al., 2007) and it can also be considered a plausible candidate for local ADAMTS-4 activation in our study.

### A critical view on the role of astrocytes

Astrocytes are additional players in dopaminergic signaling. Culture studies provided strong evidence that D1Rs and D5 receptors are expressed in rat astrocytes (Hösli and Hösli, 1986; Zanassi et al., 1999; Brito et al., 2004; Miyazaki et al., 2004). Interestingly, the D1-like DA receptor agonist SKF81297 increased intracellular cAMP levels in astrocytes (Vermeulen et al., 1994; Zanassi et al., 1999; Requardt et al., 2010). So, it is tempting to speculate that both neuronal and astrocytic D1-like DA receptors may contribute to the local increase in extracellular active proteases.

In our shRNA experiments, the knockdown of ADAMTS-4 and -5 was predominantly achieved in neurons, although we observed ADAMTS-4 expression in astrocytes (Supplementary Figure S3 which is in line with a previous study showing ADAMTS-4 to be expressed and to cleave BC in neuronal as well as in astrocytic cultures (Hamel et al., 2005). Thus, remaining BC cleavage activity in our knockdown experiments may be due to residual enzymatic activity of astrocytic-derived ADAMTS-4. Noteworthy, astrocytes are a rich source of BC as well (Zhang et al., 2014).

### Loss of full-length lecticans vs. occurrence of fragments at synapses

Several functional consequences of the observed increase of cleaved lecticans at excitatory synapses are expected: limited disintegration of the perisynaptic ECM will support receptor mobility, volume transmission of neuromodulators, and structural changes in the synaptic architecture, as discussed above. Recent work suggests that BC controls AMPAR and K_v_1 channel clustering on parvalbumin-positive interneurons (Favuzzi et al., 2017), and our own preliminary data show a regulation of small-conductance Ca^2+^-activated K^+^ (SK2) channels by BC in principal cells (Song et al., 2016), raising the possibility that proteolytic degradation may thus locally change excitability and efficacy of synapses. At the same time, the lectican fragments may exert additional signaling functions. The new and emerging concept of matricryptins as biologically active ECM fragments containing a cryptic site normally not exposed in the intact molecule (Davis et al., 2000), which was developed for non-neuronal tissues, may also apply to the neural ECM. This view is supported by the fact that the N-terminal BC fragment was shown to bind to HA and fibronectin, leading to enhanced cell adhesion in glioma (Hu et al., 2008).

The decline in cleaved BC around synapses already 10 min after stimulation of intracellular cAMP levels (Figure 7B) could point to limited retention of the fragment at perisynaptic sites where it might act as a bioactive molecule transiently inducing downstream signaling cascades.

### Conclusion: Importance of DA-modulated ECM remodeling for physiological and pathological plasticity

Homeostatic and Hebbian plasticity mechanisms interact to keep neural circuits healthy and adaptable. We have shown that induction of homeostatic plasticity in neuronal cell cultures also increased BC processing at both excitatory and inhibitory synapses (Valenzuela et al., 2014), arguing for a plasticity type-specific signature of local ECM disintegration.

Under pathological conditions disturbance of plasticity often goes along with disturbances or malformation of the neural ECM [for review see (Soleman et al., 2013); (Berezin et al., 2014)]. These pathological alterations might be related to changes in dopaminergic signaling as evidenced, for instance, by deposition of CSPGs in the Lewy bodies in Parkinson patients suffering from a loss of DA neurons (DeWitt et al., 1994). This is in line with a more recent proteomic study showing massive upregulation of the hyaluronan and proteoglycan binding link protein 2 (HAPLN2) in the substantia nigra of these patients (Liu et al., 2015).

BC and its cleavage fragments are not only important for healthy brain function but also in malignant glioma, the most common and deadly primary brain tumor (Held-Feindt et al., 2006). Up-regulation of BC and its predominantly ADAMTS-derived fragments has been shown to promote glioma invasion, while uncleavable BC was unable to enhance the invasive potential and tumor progression (Viapiano et al., 2008). Evidence for a link between BC proteolysis and synapses comes from the kainate model of epilepsy, where elevation in this ADAMTS-cleaved BC fragment was shown (Yuan et al., 2002). They observed BC proteolysis in the dentate gyrus outer molecular layer is associated with diminished synaptic density, and thus it is plausible that it is involved in the kainate-induced synaptic loss and/or reorganization. Moreover, BC was shown to be implicated in drug addiction and craving. In heroin-addicted rats a clear decrease of full length BC in synaptic protein fractions of the PFC was noted (Van den Oever et al., 2010), and Lubbers et al. (2016) demonstrated that heterozygous BC mice expressing approx. 50% of the BC levels of wild-type mice, showed a progressive increase in cocaine-associated context preference over time, which may as well involve an interplay between ECM and dopaminergic modulation. Altogether, these findings highlight that fine-tuned and locally controlled proteolytic processing of mature ECM components may have important functional implications also for the diseased or addicted brain, potentially making it an interesting target for new treatment strategies. These findings highlight that fine-tuned and locally controlled proteolytic processing of mature ECM components has important functional implications for the brain.

## Acknowledgement

The authors gratefully acknowledge expert technical assistance by Kathrin Hartung and Katrin Böhm. This work was funded by the Deutsche Forschungsgemeinschaft (SFB779 TPB14 to A.D., R.F. and C.S.; SFB779 TPB16N to M.F.K.H., SFB779 TPB09 to E.G.). Work in the authors’ labs was supported by funding from the EU (Marie Curie ITN ECMED to A.D., R.F., E.G. and C.S).

## Author’s contribution

A.D., R.F. and C.S. designed the study, supervised data analysis and edited the manuscript. R.K. designed, generated and tested the AAVs used for ADAMTS knockdown. J.M. performed all in vitro experiments and wrote the manuscript. H.N. and M.F.K.H. performed the in vivo experiment and M.F.K.H. contributed to the manuscript. C.G. provided the bPAC construct and helped with the manuscript. A.B. collaborated in the brain fractionations and statistics. E.G. discussed the project and edited the manuscript.

