## supplementary figures for "Dopamine modulates the integrity of the perisynaptic extracellular matrix at excitatory synapses"

**Supplementary Figure S1: D1R-positive puncta can be found in close vicinity to GAD65 and D2Rs are expressed in dissociated rat cortical cultures**

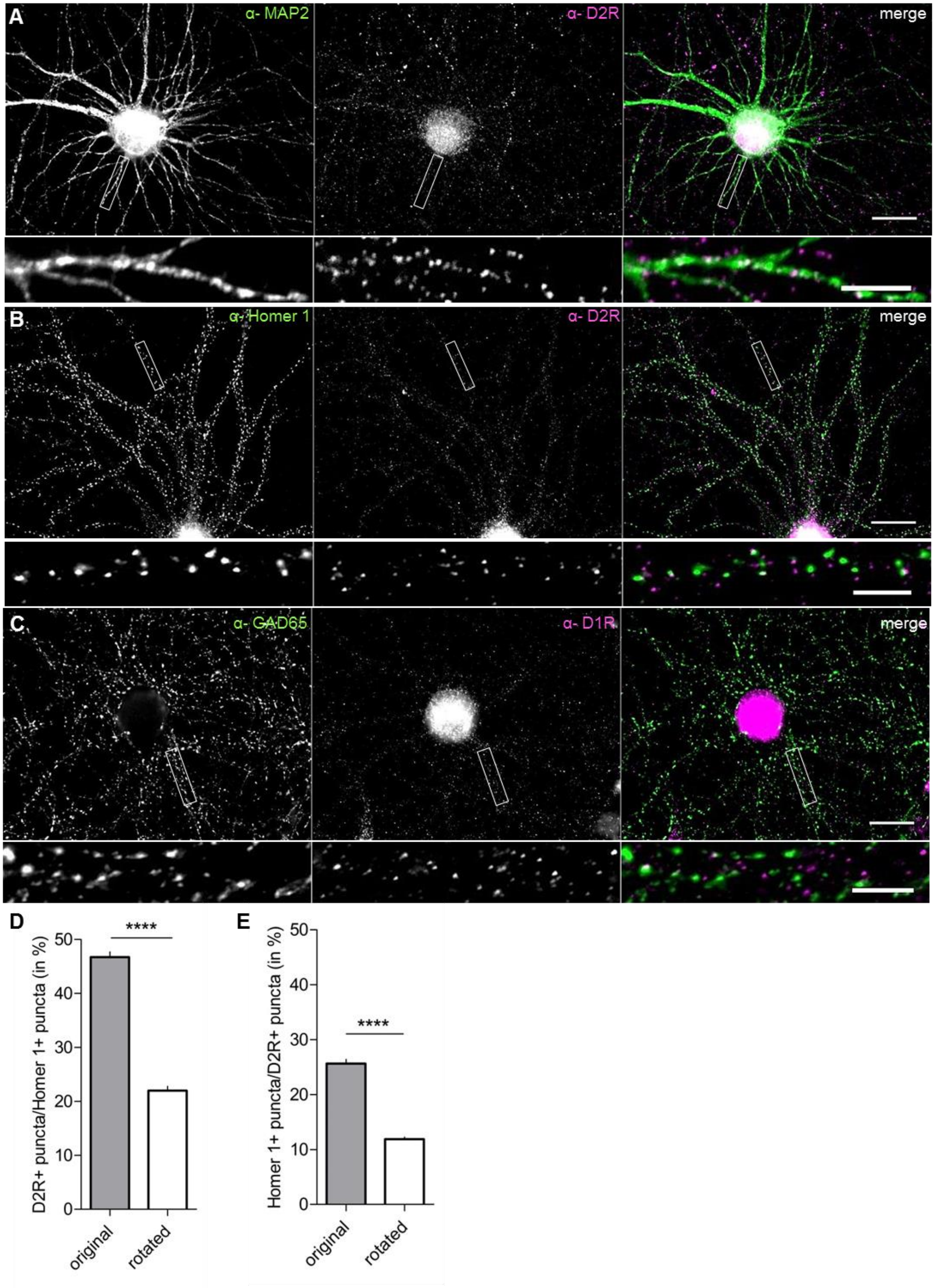

**Supplementary Figure S1: D1R-positive puncta can be found in close vicinity to GAD65 and D2Rs are expressed in dissociated rat cortical cultures**

**A)** Rat dissociated cortical neurons (DIV21) were stained for the dendritic marker MAP2 (green) and D2R (magenta). D2Rs appear as bright puncta along dendrites (scale bar: 20 µm; close-up: 5 µm). **B)** D2Rs (magenta) are found in close vicinity of Homer 1-positive excitatory synapses (green) on rat dissociated cortical neurons (DIV21) (scale bar: 20 µm; close-up: 5 µm). **C)** Rat dissociated cortical neurons (DIV21) were stained for the inhibitory synaptic marker GAD65 (green) and D1Rs (magenta). D1R-positive puncta are in close vicinity of GAD65-positive synaptic puncta (scale bar: 20 µm; close-up: 5 µm). **D)** Around 47 % of D2R-positive puncta are in close vicinity of Homer 1-positive synaptic puncta. The analysis of a 90° rotated image served as a quality control (original, 46.75 ± 0.9586, n = 8; rotated, 21.99 ± 0.8152, n = 8; average % ± SEM %; Paired t test; **** $P < 0.0001$). **E)** Around 25 % of Homer 1-positive synaptic puncta are in close vicinity of D2R-positive puncta. The analysis of a 90° rotated image served as a quality control (original, 25.66 ± 0.7576, n = 8; rotated, 11.87 ± 0.3838, n = 6; average % ± SEM %; Paired t test; **** $P < 0.0001$).

**Supplementary Figure S2: Validation of knockdown efficiency of shRNA constructs shA4 and shA5**

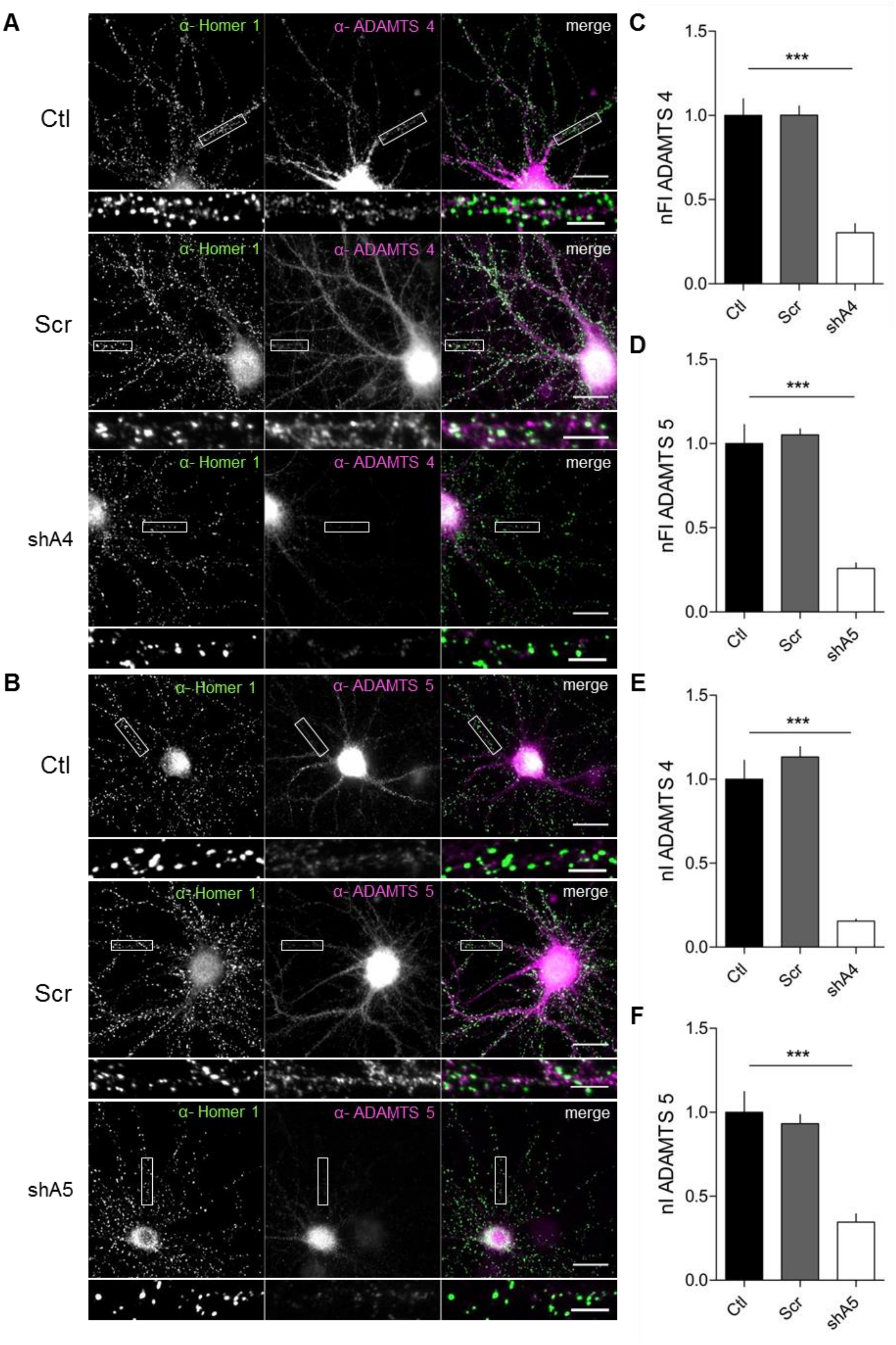

**Supplementary Figure S2: Validation of knockdown efficiency of shRNA constructs shA4 and shA5**
**A)** Rat dissociated cortical cultures (DIV14) were either non-infected (Ctl) or infected with scramble (Scr) or shADAMTS 4.2 (shA4). At DIV21 cultures were stained for the synaptic marker Homer 1 (green) and ADAMTS 4 (magenta) (scale bar: 20 µm; close-up: 5 µm). **B)** Rat dissociated cortical cultures (DIV14) were either non-infected (Ctl) or infected with Scr or shADAMTS 5.2 (shA5). At DIV21 cultures were stained for the synaptic marker Homer 1 (green) and ADAMTS 5 (magenta) (scale bar: 20 µm; close-up: 5 µm). **C)** Validation of knockdown efficiency at Homer 1-positive synapses revealed that construct shA4.2 shows an efficient knockdown (Ctl, 1 ± 0.0995, n = 5; Scr, 1.002 ± 0.0546, n = 5; shA4, 0.3034 ± 0.0543, n = 6; average ± SEM; One-way ANOVA; P = 0.001; Dunnett's Multiple Comparison Test; *** P<0.001). **D)** Also, at Homer 1-positive synapses construct shA5.2 displays an efficient knockdown for ADAMTS 5 (Ctl, 1 ± 0.1133, n = 5; Scr, 1.052 ± 0.0352, n = 4; shA5, 0.2588 ± 0.0326, n = 4; average ± SEM; One-way ANOVA; P < 0.001; Dunnett's Multiple Comparison Test; *** P<0.001). **E,F)** Knockdown efficiency was proved using western blot analysis (**E:** Ctl, 1 ± 0.113, n = 5; Scr, 0.9759 ± 0.0246, n = 5; shA4, 0.2461 ± 0.0188, n = 3; average ± SEM; One-way ANOVA; P = 0.001; Dunnett's Multiple Comparison Test; *** P<0.001; **F:** Ctl, 1 ± 0.1234, n = 6; Scr, 0.9407 ± 0.0379, n = 6; shA5, 0.3465 ± 0.0487, n = 4; average ± SEM; One-way ANOVA; P = 0.0003; Dunnett's Multiple Comparison Test; *** P<0.001) (nFI = normalized fluorescent intensity; nI = normalized intensity)

**Supplementary Figure S3: ADAMTS 4 is expressed by neurons and astrocytes**

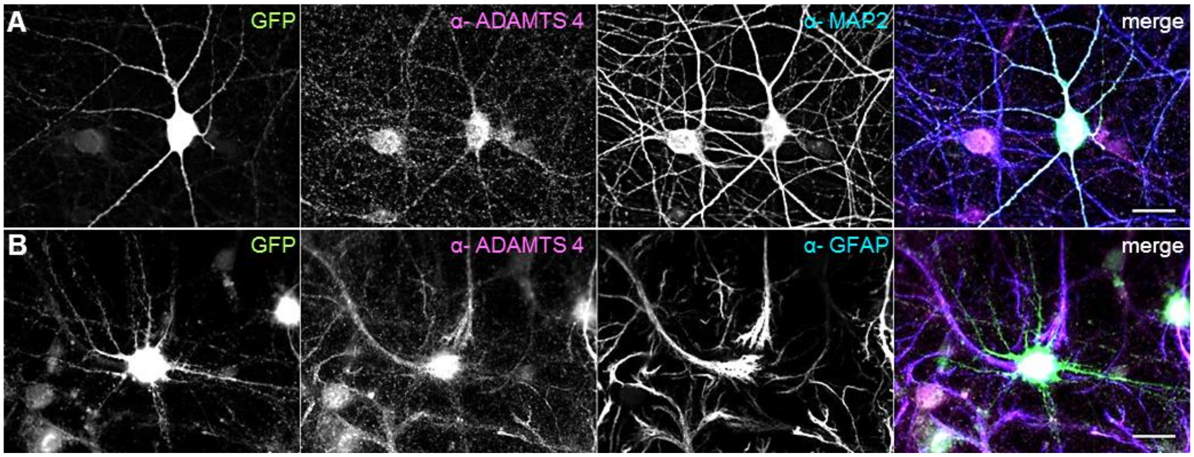


**A,B)** Rat dissociated cortical cultures (DIV14) were infected using shADAMTS4.1-AAV-GFP. At DIV 21 cells were stained for ADAMTS 4 (magenta) and the somato-dendritic marker MAP2 (A,blue) or the astrocytic marker GFAP (B, blue). The strong GFP signal (green) indicates the successful infection with the used AAV. Astrocytes display no GFP signal suggesting no infection with the used AAV (scale bar: 20 µm).
